# STAG2 loss-of-function affects short-range genomic contacts and modulates urothelial differentiation in bladder cancer cells

**DOI:** 10.1101/2020.08.06.240457

**Authors:** Laia Richart, Eleonora Lapi, Vera Pancaldi, Mirabai Cuenca, Enrique Carrillo-de-Santa Pau, Miguel Madrid-Mencía, Hélène Neyret-Kahn, François Radvanyi, Juan A. Rodríguez, Yasmina Cuartero, François Serra, François Le Dily, Alfonso Valencia, Marc A. Marti-Renom, Francisco X. Real

**Author notes:** Computational Biology Group, Precision Nutrition and Cancer Program, IMDEA Food Institute, 28049 Madrid, Spain. equal contribution (Laia Richart, Eleonora Lapi).

## Abstract

Cohesin exists in two variants, containing either STAG1 or STAG2. *STAG2* is one of the most commonly mutated genes in human cancer, and a major bladder cancer tumor suppressor. Little is known about how its inactivation contributes to tumor development. Here, we analyze the genomic distribution of STAG1 and STAG2 and perform STAG2 loss-of-function experiments using RT112 bladder cancer cells; we then analyze the resulting genomic effects by integrating gene expression and chromatin interaction data. Cohesin-STAG2 is required for DNA contacts within topological domains, but not for compartment maintenance of domain boundary integrity. Cohesin-STAG2-mediated interactions are short-ranged and engage promoters and gene bodies with higher frequency than those mediated by cohesin-STAG1. STAG2 knockdown resulted in a modest but consistent down-regulation of the luminal urothelial differentiation signature, mirroring differences between STAG2-high and STAG2-low bladder tumors. Both lost and gained contacts were enriched among STAG1/STAG2 common sites as well as STAG2-enriched sites. Contacts lost upon depletion of STAG2 were significantly assortative, indicating their proximity at the 3D level, and were associated with changes in gene expression. Overall, our findings indicate that, in urothelial cells, STAG2 is required for the establishment and/or maintenance of DNA looping that, in turn, sustains the luminal differentiation program. This mechanism may contribute to the tumor suppressor function of STAG2 in bladder cancer.

## INTRODUCTION

*STAG2* encodes a subunit of the cohesin complex and is one of the most commonly mutated genes in human cancer (Lawrence et al. 2014). Among cohesin-associated genes, it harbours the highest frequency of predicted pathogenic mutations (Romero-Pérez et al. 2019). Focal deletions on the X chromosome involving *STAG2* were first detected in glioblastomas (Solomon et al. 2011). Subsequently, massive parallel sequencing allowed the detection of point mutations in a variety of human tumors including urothelial bladder cancer (UBC) (Solomon et al. 2011; Balbás-Martínez et al. 2013; Solomon et al. 2013; Guo et al. 2013; Taylor et al. 2014), Ewing sarcoma (Brohl et al. 2014; Crompton et al. 2014), myelodysplastic syndrome, and acute myeloid leukaemia (AML) (Kon et al. 2013; Thota et al. 2014). The majority of *STAG2* mutations reported in cancer lead to a premature stop codon and the absence of protein (De Koninck and Losada 2016), but loss of expression can also result from gene deletion and/or methylation (Guo et al. 2013; Shen et al. 2016; Cheung et al. 2016). Mutations were first reported almost a decade ago and there is increasing evidence for a role of *STAG2* as a tumor suppressor; yet, the mechanisms whereby its inactivation contributes to cancer remain elusive. In UBC, *STAG2* mutations occur mainly in indolent tumors (Balbás-Martínez et al. 2013; Solomon et al. 2013; Guo et al. 2013; Taylor et al. 2014). By contrast, in Ewing sarcoma they are associated with aggressive neoplasms (Brohl et al. 2014; Crompton et al. 2014), emphasizing the need to perform functional analyses in appropriate model systems to identify potential tissue-specific effects.

The cohesin complex is composed of SMC1, SMC3, RAD21, and either STAG1 or STAG2. As a result, two versions of the complex exist in somatic cells with potentially distinct biological functions (Remeseiro and Losada 2013). In this regard, knockout mouse models have revealed that STAG1 plays a predominant role in telomeric cohesion, while STAG2 plays a more important role in cohesion at chromosome arms or in centromeric regions (Canudas and Smith 2009; Remeseiro et al. 2012a; Remeseiro et al. 2012b). The well-established role of cohesin in chromosome segregation initially suggested that *STAG2* inactivation in cancers might be associated with aneuploidy (Solomon et al. 2011). However, genetic analyses of UBC and AML strikingly showed that *STAG2*-mutant tumors were genomically stable (Balbás-Martínez et al. 2013; Welch et al. 2012), supporting the importance of additional molecular mechanisms. These findings are in contrast with a higher rate of somatic copy number changes - but not ploidy - in Ewing sarcoma (Crompton et al. 2014).

There is increasing evidence that cohesin participates in a variety of processes beyond chromosome segregation, including DNA repair and replication, chromatin organization, and gene regulation (De Koninck and Losada 2016). An increased understanding of these processes has emerged from the recent use of Chromosome Conformation Capture (3C) technologies (Dekker et al. 2002) including Hi-C (Lieberman-Aiden et al. 2009), which revealed that genomes are folded into complex, hierarchically organized, 3D structures playing a key role in essential processes (e.g. transcription). These structures span a wide range of length scales: from large chromosomal domains and compartments, ~1Mb self-interacting domains (Topologically Associating Domains or TADs), to DNA loops connecting promoters and gene regulatory elements (Rowley et al. 2018). Cohesin, together with the chromatin insulator CTCF, contributes to TAD border definition by means of loop extrusion (Sanborn et al. 2015; Guo et al. 2015; Fudenberg et al. 2016; Rao et al. 2017; Kim et al. 2019), as well as to intra-TAD promoter-enhancer interactions (Kagey et al. 2010; Schaaf et al. 2013). A large fraction of genomic sites targeted by cohesin are simultaneously bound by STAG1, STAG2, and CTCF, yet a few STAG1-only and STAG2-only sites occur in the genome (Remeseiro et al. 2012; Kojic et al. 2018; Cuadrado et al. 2019). The latter are depleted of CTCF and are enriched in enhancers and transcription factor binding sites. Knockdown experiments showed that, upon STAG2 depletion, cohesin-STAG1 does not bind to the STAG2-only sites (Kojic et al. 2018; Cuadrado et al. 2019; Casa et al. 2020; Viny et al.), suggesting that cohesin-STAG2 has unique distribution and functions whose role in tumor suppression is yet to be determined.

Despite UBC having the highest frequency of *STAG2* mutations, there are no studies on the role of STAG2 in urothelial cells at the genomic level. UBC is a heterogeneous cancer with two broad histological subtypes (Gui et al. 2011; McConkey et al. 2010): low-grade/papillary (75-80% of cases) - which tend to be non muscle-invasive - and solid/muscle-invasive (20-25% of cases). The latter can present with variable phenotypes: some tumors preserve urothelial/luminal features while others show basal/squamous characteristics. Up to 40% of papillary tumors harbor *STAG2* inactivation, which is significantly associated with activating *FGFR3* mutations and low levels of genomic instability (Balbás-Martínez et al. 2013; Taylor et al. 2014). Conversely, *STAG2* mutations occur only in 12-15% of muscle-invasive tumors. Among them, *STAG2* mutations are enriched in tumors with urothelial/luminal differentiation and *FGFR3* mutations (Kamoun et al. 2020), suggesting that they represent the invasive counterpart of a subset of papillary tumors. Several questions arise from these clinical-molecular associations, including the mechanistic basis of the association with specific transcriptomic signatures and urothelial differentiation.

Here, we set out to explore the contribution of STAG2 to genome organization and the urothelial transcriptional program in RT112, a well-characterized bladder cancer cell line displaying luminal features, mutant *FGFR3* and wild-type *STAG2* (Earl et al. 2015). Using Hi-C in combination with ChIP-Seq for cohesin subunits, and RNA-Seq, we show that the cohesin-STAG2 complex is important for the formation and/or maintenance of DNA contacts within TADs, but not for the integrity of their boundaries. Chromosomal interactions mediated by cohesin-STAG2 are short-ranged and engage promoters and gene bodies with higher frequency than those mediated by cohesin-STAG1, in agreement with notions of compartmentalization in the molecular processes undertaken by the two types of cohesin complexes (Canudas and Smith 2009; Remeseiro et al. 2012a; Remeseiro et al. 2012b; Kojic et al. 2018; Casa et al. 2020; Viny et al.). Depletion of STAG2 leads to rewiring of short-range contacts and concomitant changes in the expression of selected luminal/basal signature genes.

## RESULTS

### STAG2-enriched cohesin localizes to active enhancers and promoters independently of CTCF

We profiled the genome-wide distribution of STAG2 and STAG1 in RT112 cells by chromatin immunoprecipitation followed by deep sequencing (ChIP-Seq). SMC1, common to both cohesin complexes, was used as control. Three categories of cohesin-bound genomic positions were identified based on differential binding of STAG2 and STAG1: common (n=35,321), STAG1-enriched (n=5,007), and STAG2-enriched (n=2,330) (Figure 1A and Supplementary Figure 1A). Common positions were occupied by either complex variant and showed comparably high read density for both STAG1 and STAG2. STAG1-enriched positions showed higher read density for STAG1 than for STAG2. In contrast, STAG2-enriched positions had higher STAG2 read density than STAG1, but showed the lowest read density of all categories. SMC1 was present in the three categories of cohesin-bound positions (Figure 1A and Supplementary Figure 1A). Peak-centered ChIP-Seq read density plots revealed a pattern of sharp and narrow peaks for STAG1 and STAG2 around common and STAG1-enriched cohesin positions. STAG2 peaks at STAG2-enriched sites were broader (Figure 1B), suggesting higher cell-to-cell variability or greater dynamics of this complex variant (Kojic et al. 2018).

**Figure 1.**
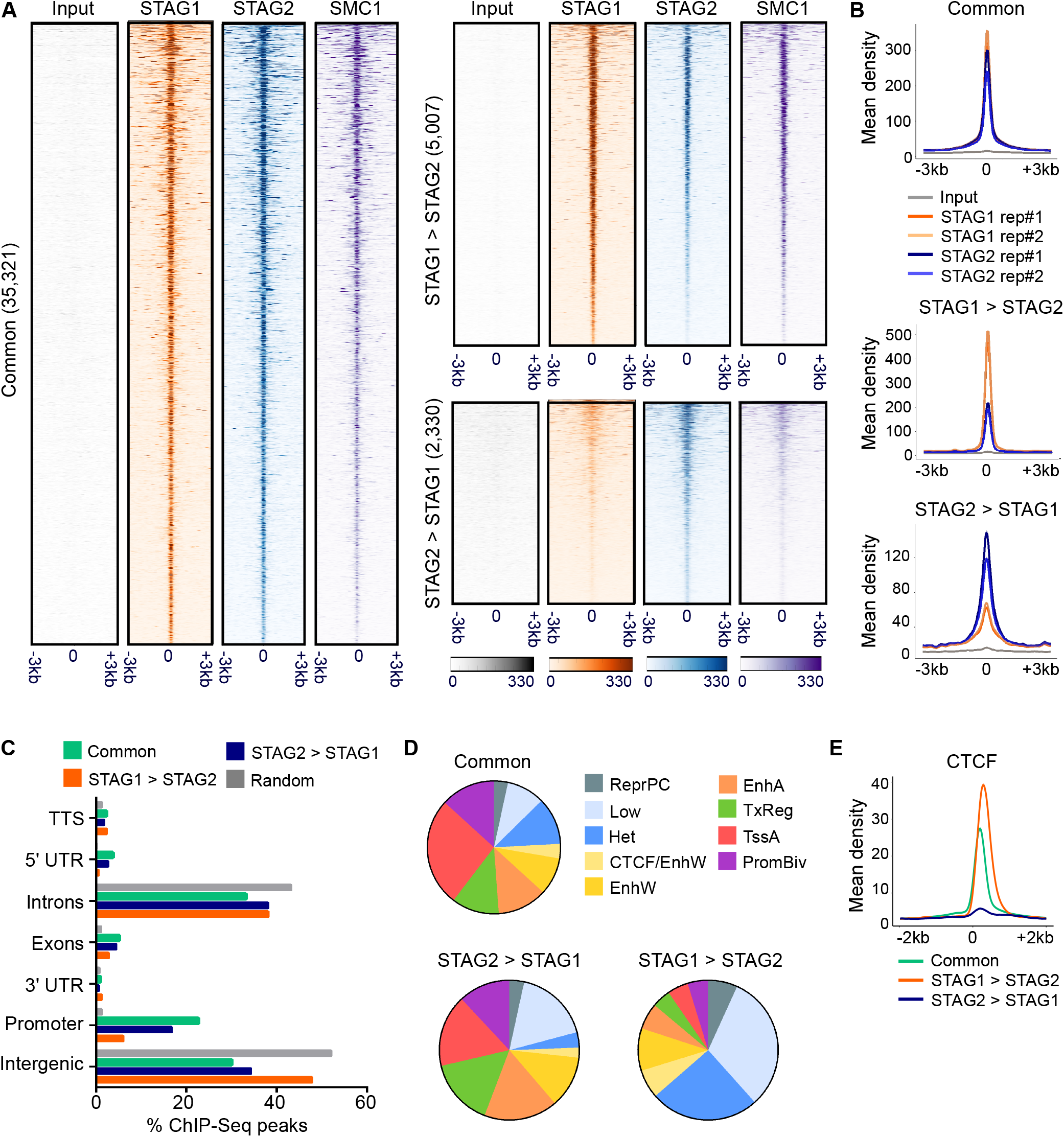
STAG1 and STAG2 show both overlapping and unique distributions over genomic elements and chromatin states in RT112 cells. (A) ChIP-Seq read density heatmaps for STAG1, STAG2, and SMC1 at common, STAG1-enriched (STAG1 > STAG2), and STAG2-enriched (STAG2 > STAG1) cohesin positions within a peak-centered 6kb window. (B) Read density distribution for STAG1 and STAG2 at common, STAG1-enriched, and STAG2-enriched positions within a peak-centered 6kb window. (C) Bar-plot diagram showing the distribution of common, STAG1-enriched, and STAG2-enriched cohesin positions over genomic elements. (D) Distribution of cohesin-bound genomic sites throughout chromatin states identified in RT112 cells by ChromHMM and based on combinations of histone modifications and CTCF (see Supplementary Figure 1C for definition of chromatin states). (E) Peak-centered enrichment plot for CTCF over the three categories of cohesin-bound positions showing relative depletion in STAG2-enriched sites.

Analysis of enrichment over genomic elements revealed no differences in the distribution of STAG1 and STAG2 when considered independently (Supplementary Figure 1B). STAG1-enriched positions were comparatively more abundant in intergenic regions whereas common and STAG2-enriched peaks showed higher overlap with promoters, exons, and 5’ Untranslated Regions (5’ UTR) (Figure 1C and Supplementary Figure 1B). We then investigated cohesin enrichment over 9 chromatin states defined by combinations of histone modifications and CTCF in RT112 cells (Supplementary Figure 1C). Similarly to the overlap with genomic features, distribution of STAG1 and STAG2 according to chromatin states was highly comparable unless their relative enrichment was accounted for (Figure 1D and Supplementary Figure 1D). STAG1-enriched positions characteristically overlapped with transcriptionally inactive chromatin marked by H3K27me3 (“ReprPC”), H3K9me3 (“Het”), or low levels of the assessed histone modifications (“Low”), in addition to chromatin domains bound by CTCF (“CTCF/EnhW”). Conversely, both common and STAG2-enriched sites were mostly distributed over transcriptionally active genes, promoters, and enhancers (Figure 1D). Peak-centered density plots highlight the differential dependency of the three categories of cohesin-bound genomic positions on CTCF (Figure 1E). Consistently, motif analysis showed that STAG1-enriched positions were significantly enriched for the CTCF binding motif, whereas STAG2-enriched positions displayed binding motifs of transcription factors participating in cancer, including ASCL1, MYCN, and KLF5 (Supplementary Figure 1E) (Du et al. 2016; Qin et al. 2015; Sabari et al. 2017). These findings are largely in agreement with previous observations in other cell types (Kojic et al. 2018; Cuadrado et al. 2019). Unlike in mES cells (Cuadrado et al. 2019), we did not detect higher overlap with H3K27me3 domains among STAG2-enriched sites (Figure 1D).

### Chromosomal compartments and TAD boundaries are resilient to STAG2 depletion

To assess the contribution of STAG2 to chromatin architecture and transcriptional regulation in RT112 cells, we efficiently silenced STAG2 with two shRNAs (sh1 and sh2) - using a non-targeting shRNA as control (shNT) (Figure 2A) - and performed Hi-C and RNA-Seq experiments. Given the role of cohesin in sister chromatid segregation (Morales and Losada 2018), we first assessed whether STAG2 depletion affected ploidy. Visual exploration of the number of reads per bin in genome-wide 100kb contact matrices did not unveil gross genomic differences in ploidy between control and STAG2-depleted cells (Supplementary Figure 2).

**Figure 2.**
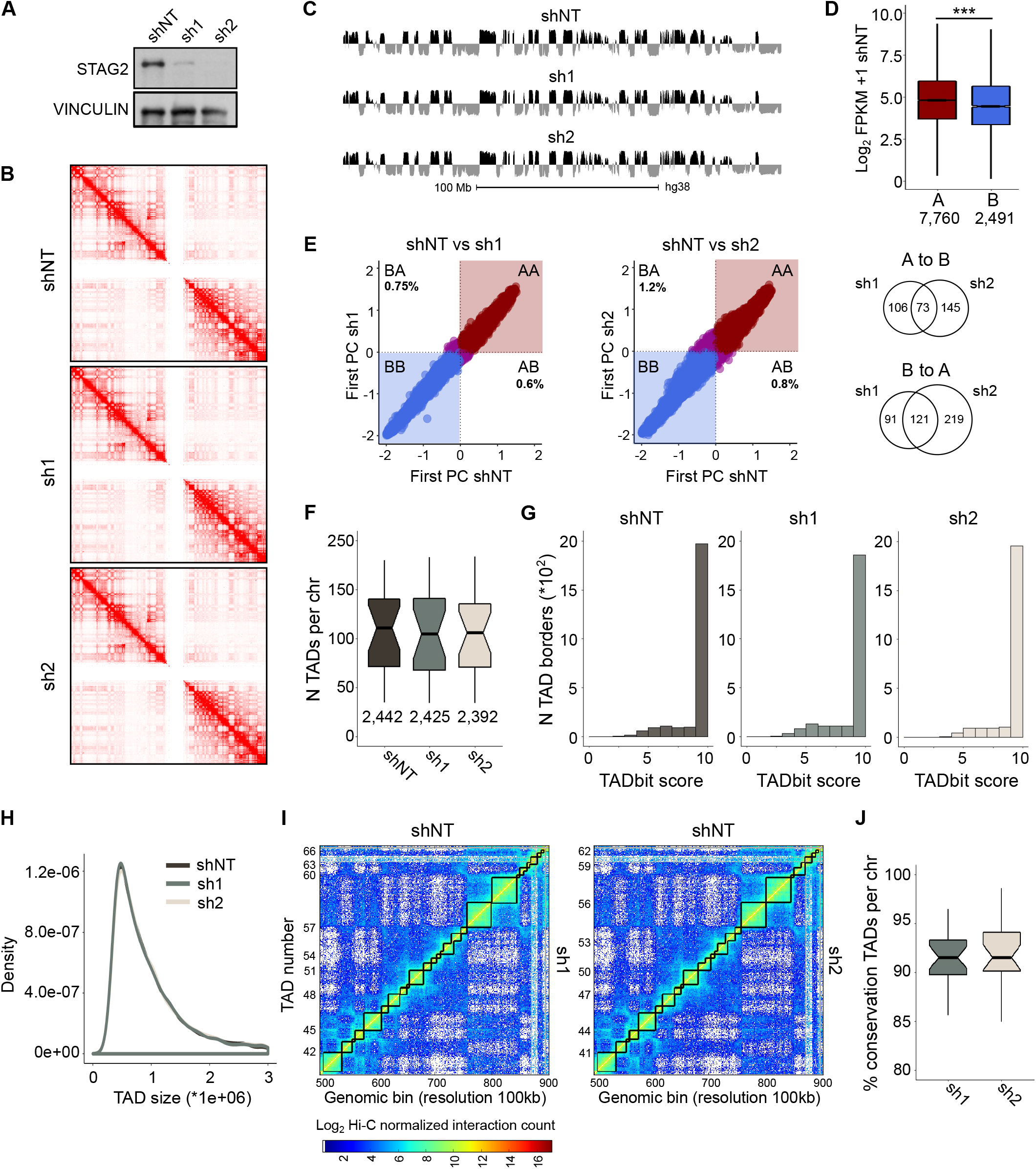
Down-regulation of STAG2 in RT112 cells does not interfere with A/B compartments or TAD boundaries. (A) Western blot analysis of control (shNT) and STAG2-silenced RT112 cells showing efficient depletion of STAG2 at the protein level. (B) Hi-C matrices for chr2 at 500kb resolution in cells transduced with control or STAG2-targeting shRNAs. The darker red reflects a greater frequency of interaction. (C) Compartment tracks for chr2 at 100kb resolution as determined by the values of the first principal component (PC1) in control and STAG2-silenced cells. (D) Expression, as defined by RNA-Seq (Log2 FPKM), of genes within compartments A and B. As expected, genes assigned to compartment A are more transcriptionally active than genes in compartment B. t-test: ***, *P* < 0.001. (E) Scatterplot of PC1 values of the eigenvectors of intrachromosomal interaction matrices for control and STAG2-silenced cells. The Venn diagrams show the overlap in terms of compartment-switching bins between sh1 and sh2. (F) Effect of STAG2-depletion on the number of TADs per chromosome. (G) Histograms depicting the strength of the TAD borders detected in control and STAG2-silenced cells, according to the TADbit score. (H) Density plot depicting the distribution of TAD sizes identified in control and STAG2-silenced cells. (I) Hi-C normalized interaction matrices for chr2 at 100kb resolution comparing TAD organization in control and STAG2-silenced cells. (J) Effect of STAG2-depletion on the conservation of TAD borders. Boxplot notches represent the confidence interval around the median.

To determine whether STAG2 loss resulted in major changes in chromatin organization, we first explored the effect on genomic compartments. The genome is segregated into two major compartments, named A and B, that differ in their gene density, epigenetic modifications, and transcriptional output. Overall, the A compartment contains transcribed genes and active histone modifications while the B compartment generally encompasses inactive genes with histone modifications associated with a repressed transcriptional state (Rowley et al. 2018). A and B compartments were defined using the first principal component (PC) obtained by eigenvector decomposition of normalized Hi-C matrices at 100kb resolution (Figure 2B) in combination with information on gene density and transcriptional activity (Figure 2C). In control cells, genomic regions assigned to compartment A were comparatively gene-rich (7,760 genes in A vs. 2,491 genes in B) and genes therein were expressed at significantly higher levels (Figure 2D). Segregation into A/B compartments in shNT and shSTAG2 RT112 cells was highly correlated, with 45.3% of the genome consisting of constitutive A-type domains and 54.7% classified as B-type (Figure 2E). In agreement with the preferential association of cohesin with genes and their regulatory elements, most of its binding sites were found in compartment A, and no major differences were observed between the three categories of cohesin-bound genomic sites (common: 67%, STAG1 enriched: 66%, STAG2 enriched: 67%). Upon STAG2 knock-down, only 0.6-0.8% and 0.75-1.2% of the genome underwent A-to-B and B-to-A compartment changes, respectively (Figure 2E). A significant degree of overlap was detected between STAG2-silenced cells in terms of genes “flipping” compartments, suggesting small but consistent effects (Figure 2E).

We then explored the possibility that STAG2 depletion may interfere with the organization of the genome into TADs. Using normalized 100kb contact maps, we identified a total of 2,442, 2,425, and 2,392 TAD borders of comparable strength (TADbit score > 5) in shNT, sh1, and sh2 cells, respectively (Figure 2F,G). The number and size of TADs was similar among conditions (Figure 2F,H) and, as revealed by alignment of their boundaries, they were highly conserved (average conservation with respect to shNT: sh1 90.8%, sh2 91.5%). The small decrease in TAD number in STAG2-depleted cells (Figure 2H,I) might result from merging of adjacent TADs. In agreement with previous work (Kojic et al. 2018; Cuadrado et al. 2019; Casa et al. 2020), our results indicate that megabase-scale architectural compartments and TAD borders are resilient to reduced STAG2 protein levels.

### STAG2-enriched cohesin mediates short range intra-TAD interactions

Cohesin contributes to the 3D conformation of chromatin at the submegabase scale in a cell-type specific fashion through both long-range constitutive interactions and short-range promoter-enhancer contacts that regulate transcription (Phillips-Cremins et al. 2013). STAG2 knock-down resulted in modest, statistically significant (FDR < 0.05), changes in expression levels of a subset of genes (sh1, n=510; sh2, n= 438), with a similar number of up- and down-regulated transcripts (Supplementary Figure 3A). Gene expression changes induced by either shRNA were positively correlated and 20-32% of significantly up- and down-regulated genes were common to both shRNAs (Supplementary Figure 3B). Conditional *Stag2* deletion in the mouse blood compartment is associated with transcriptional dysregulation and impaired differentiation of hematopoietic stem cells (Viny et al.). We thus explored the effects of depleting STAG2 on RT112 differentiation by assessing the differential expression of gene signatures characteristic of muscle-invasive UBC molecular subtypes (Kamoun et al. 2020). GSEA (Gene Set Enrichment Analysis) revealed consistent and significant up-regulation of genes linked to the basal/squamous class, and a significant down-regulation of genes linked to the luminal papillary class in cells transduced with both STAG2-targeting shRNAs (Supplementary Figure 3C). These findings suggest that STAG2 loss leads to defective maintenance of the luminal differentiation transcriptional program (Supplementary Figure 3C). GSEA further revealed a significant overlap between genes down-regulated in STAG2-silenced RT112 cells and genes differentially expressed in “*STAG2* low” (bottom quartile) versus “*STAG2* high” (top quartile) samples from the UROMOL study, involving 476 cases of non muscle-invasive bladder cancer (Hedegaard et al. 2016) (Supplementary Figure 3D-G), thus validating the relevance of our *in vitro* system.

We hypothesized that STAG2 knockdown might affect transcription by interfering with the formation of chromatin loops engaging specific promoters and their regulatory regions. To address this question, we investigated significant contacts predicted by HOMER (FDR < 0.1) at mid resolution (20kb) in control and STAG2-depleted cells (Figure 3A). Upon STAG2 knock-down, interaction frequency, or the number of interactions between pairs of genomic regions, was significantly lower (mean interaction reads - shNT: 28.5; sh1: 24.5; sh2: 27.2) (Figure 3B). Furthermore, we found an increase in the genomic distance spanned by the interactions resulting from a loss of short-range (<250kb) and a concomitant increase in long-range contacts (>1Mb) in the two STAG2-depleted conditions (Figure 3C). These observations are largely in agreement with previous Hi-C analyses of mouse and human cells silenced for STAG2 (Kojic et al. 2018; Cuadrado et al. 2019; Viny et al.).

**Figure 3.**
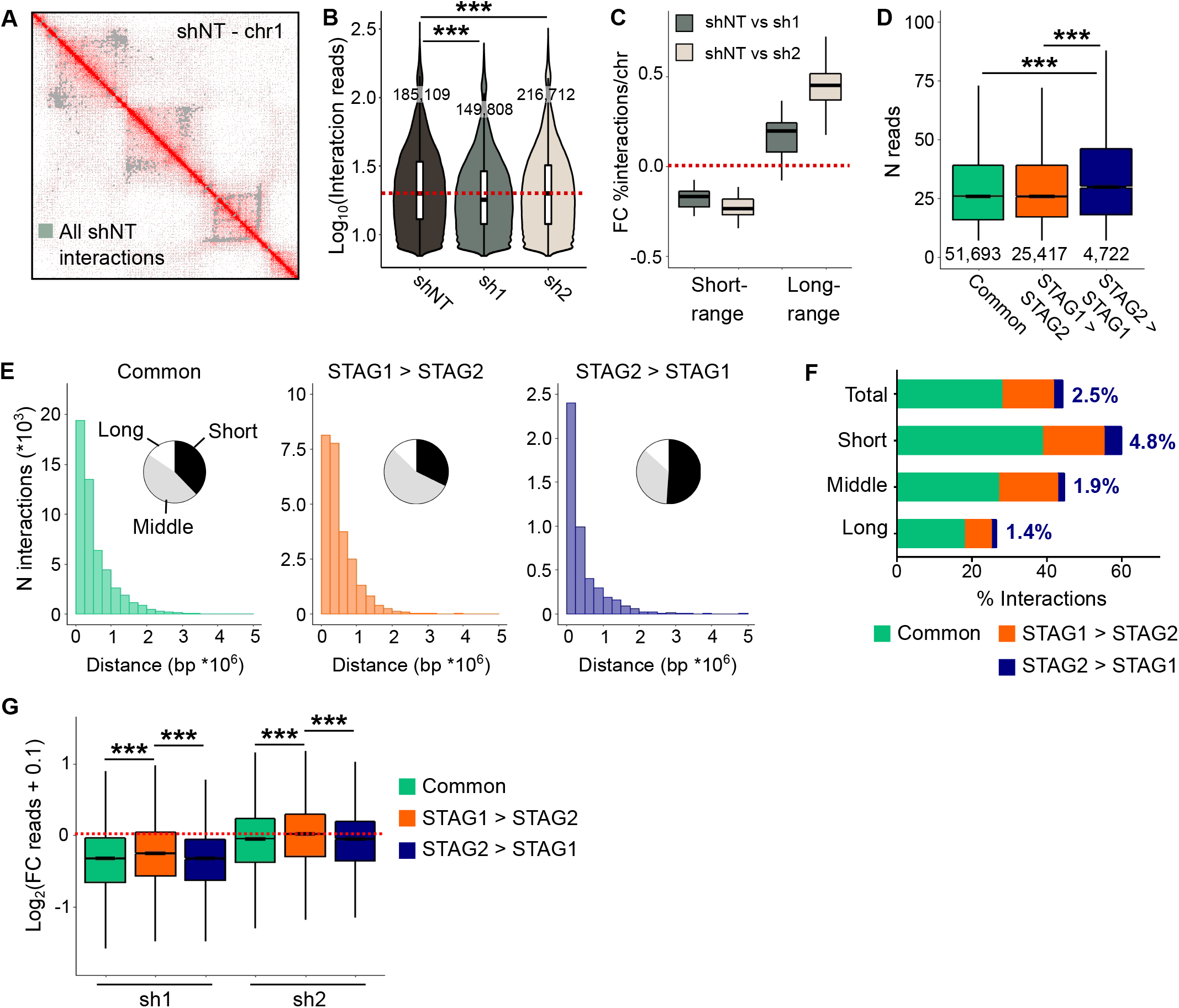
STAG2-enriched cohesin mediates short-range contacts in RT112 cells. (A) Snapshot of a normalized contact matrix of chr1 at 20kb resolution in control cells as visualized in Juicebox (Durand et al. 2016). Grey dots indicate the position of significant interactions (FDR < 0.1). The darker red reflects a greater frequency of interaction. (B) Number of reads per interaction for significant interactions (FDR < 0.1) called in control (shNT) and STAG2-depleted RT112 cells. (C) Fold change of the proportion of interactions per chromosome comparing control and STAG2-silenced cells, classified as short (< 250kb) or long-range (>1 Mb). (D) Number of reads per interaction of interactions overlapping cohesin binding sites in control cells. (E) Histograms showing the distribution of distances between the peaks of interactions in control cells overlapping common, STAG1-enriched, or STAG2-enriched cohesin genomic binding sites. The pie charts plot the proportion of short (< 250kb), mid (250kb-1Mb), and long-range (>1 Mb) interactions. STAG2-enriched overlapping interactions are distinctively short-ranged. (F) Proportion of short, middle, and long interactions overlapping the three subsets of cohesin binding sites. Numbers within the plot show the proportion of STAG2-enriched overlapping interactions. (G) Fold-change in the number of reads of contacts overlapping common, STAG1-enriched, or STAG2-enriched cohesin binding sites upon STAG2 silencing. t-test: ***, *P* < 0.001.

To dissect the effect on loop formation, we overlapped interactions in control cells with the three types of cohesin-binding sites identified in our ChIP-Seq experiments. Interactions overlapping STAG2-enriched binding sites displayed higher interaction frequencies (Figure 3D) and spanned shorter genomic distances (Figure 3E) than those engaging STAG1-enriched or common sites. Classifying control interactions according to distance [short (<250kb), mid (250kb-1Mb), long-ranged (>1Mb)] confirmed that interactions overlapping STAG2-enriched positions are more abundant among short-range contacts (Figure 3F). Importantly, upon STAG2-depletion, loss of interaction reads was higher at binding sites occupied by STAG2 (common or STAG2-enriched) (Figure 3G), supporting the specificity of the effects.

STAG2 knockdown led to a statistically significant decrease in the frequency of contacts defined in control cells (Figure 4A). To better evaluate the functional consequences of changes in DNA looping upon STAG2 silencing, we defined a collection of interactions significantly “lost” and “gained” in both shRNAs (*P* < 0.05) (Figure 4B). Lost and gained interactions showed similar distributions over A/B compartments and genomic elements (Supplementary Figures 4A,B), but differed in terms of distance between interaction peaks. While both sets of contacts were restricted to a 1Mb window, lost interactions spanned shorter distances (<250kb), reminiscent of the ones overlapping STAG2-enriched cohesin positions (Figure 4C,D and Supplementary 4C). Motif analysis of genomic regions engaged by control (shNT), lost, and gained interactions revealed that CTCF was among the top three scoring motifs enriched in control and gained interactions but not among those lost (Figure 4E). These changes echo the poor overlap between STAG2-enriched cohesin sites and CTCF observed in the ChIP-Seq analysis (Figure 1D,E and Supplementary Figures 1A,E). Importantly, the proportion of lost and gained contacts was higher among common and STAG2-enriched cohesin-overlapping interactions (Figure 4F), supporting that these changes are causally linked to silencing of STAG2.

**Figure 4.**
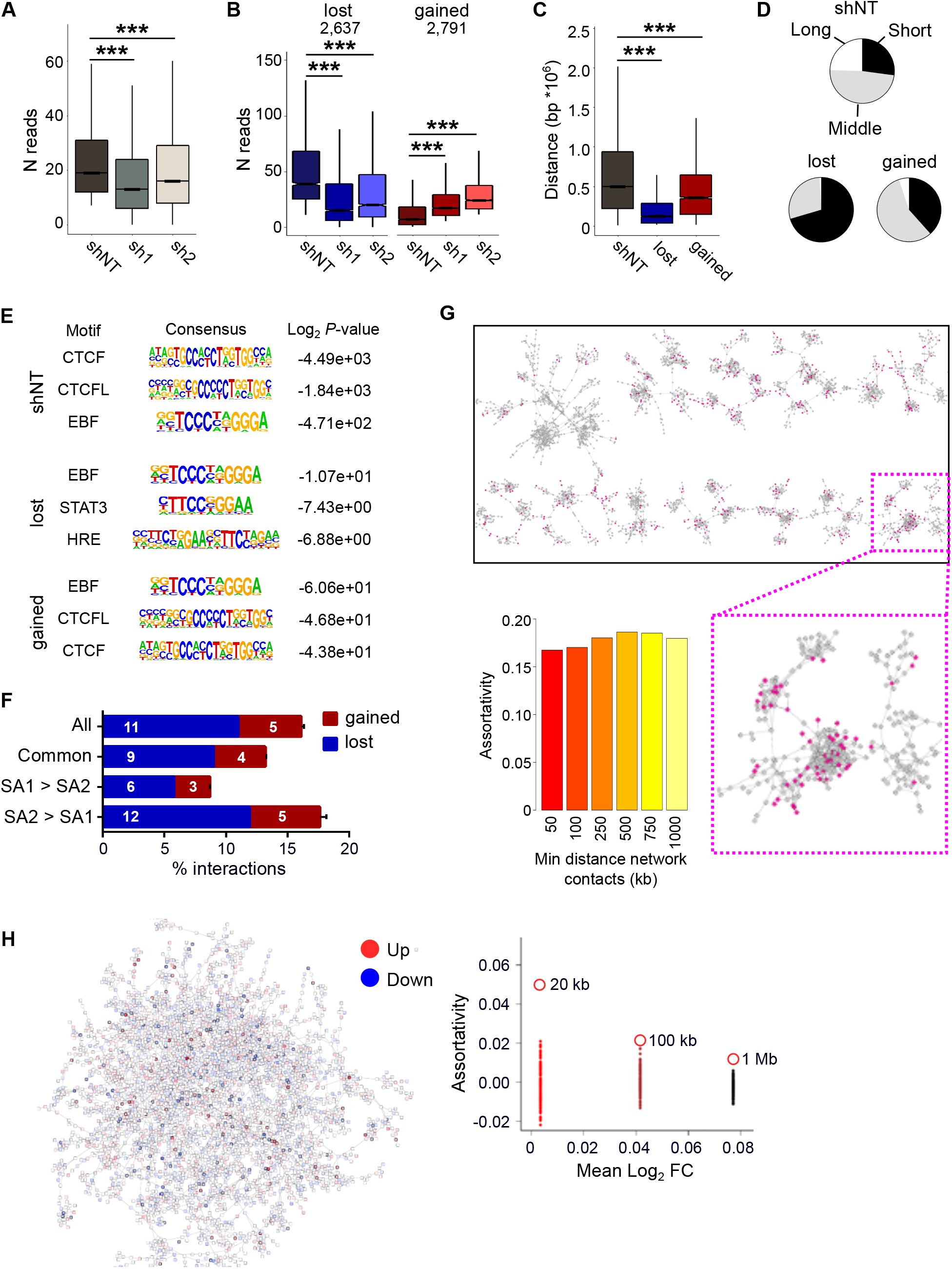
STAG2 silencing is accompanied by loss of short-ranged, assortative, contacts that do not overlap with CTCF binding sites. (A) Number of reads aligning to genomic sites engaged in interactions in control shNT cells (FDR<0.1) and in STAG2-silenced cells. (B) Number of reads of interactions significantly lost and gained. (C) Distance between peaks of control (shNT), lost, and gained interactions. (D) Proportion of short (<250kb), mid (250kb-1Mb), and long-ranged (>1Mb) contacts among control (shNT), lost, and gained interactions. (E) Motif analysis of the subsets of interactions defined in B. (F) Proportion of lost and gained interactions among control or cohesin-overlapping contacts. (G) Top: chromatin contact network generated from the 20kb resolution Hi-C interaction map of control cells, showing in pink the nodes involved in contacts that are lost upon STAG2 silencing. Bottom: chromatin assortativity of nodes that lose contacts as the network is filtered eliminating contacts spanning short distances. (H) Left: Network of gene contacts inferred from 20kb resolution Hi-C interaction matrices of control cells (including only genes for which expression values could be calculated) showing log2 fold change of expression between STAG2-silenced and control cells. Nodes in red are up-regulated in STAG2-silenced cells while nodes in blue are down-regulated. Significantly regulated genes are shown with black borders. Right: Assortativity of fold-change of expression values between STAG2-silenced and control cells measured on a network of gene-gene 3D interactions, calculated using networks generated by different binnings of Hi-C data (large empty red circles 20kb, 100kb and 1Mb), compared to random expectations for assortativity values (small filled circles). t-test: ***, *P* < 0.001.

### Differential assortativity of lost and gained interactions

To assess whether the loci that lose or gain interactions upon STAG2 silencing are located close to each other in 3D, we performed a chromatin assortativity analysis whereby the genome is represented as a network of interacting nodes and each node is a chromatin fragment (Pancaldi et al. 2016). Loci involved in lost interactions were more assortative than expected by chance, suggesting that they tend to be close to each other in the 3D space. Assortativity of these regions was stable on networks in which contacts spanning shorter distances were eliminated, reaching a maximum for interactions that span less than a megabase (intra-TAD) (Figure 4G).

To investigate whether genes that are differentially regulated in the same direction in STAG2 depleted cells are closer to each other in 3D than expected by chance, we estimated the assortativity of changes in gene expression in STAG2-silenced vs. control cells. Interestingly, we found high assortativity only when considering Hi-C interactions binned at 20kb resolution but not for lower resolution networks (>100kb) (Figure 4H). These observations are consistent with the notion that the vast majority of lost interactions were short-range (<250kb) (Supplementary Figure 4C) and that lost contacts potentially affected gene expression (Figure 5C,D). On the contrary, gained interactions did not show any patterns of assortativity, suggesting a lack of a functional consistency between loci affected by gained interactions.

**Figure 5.**
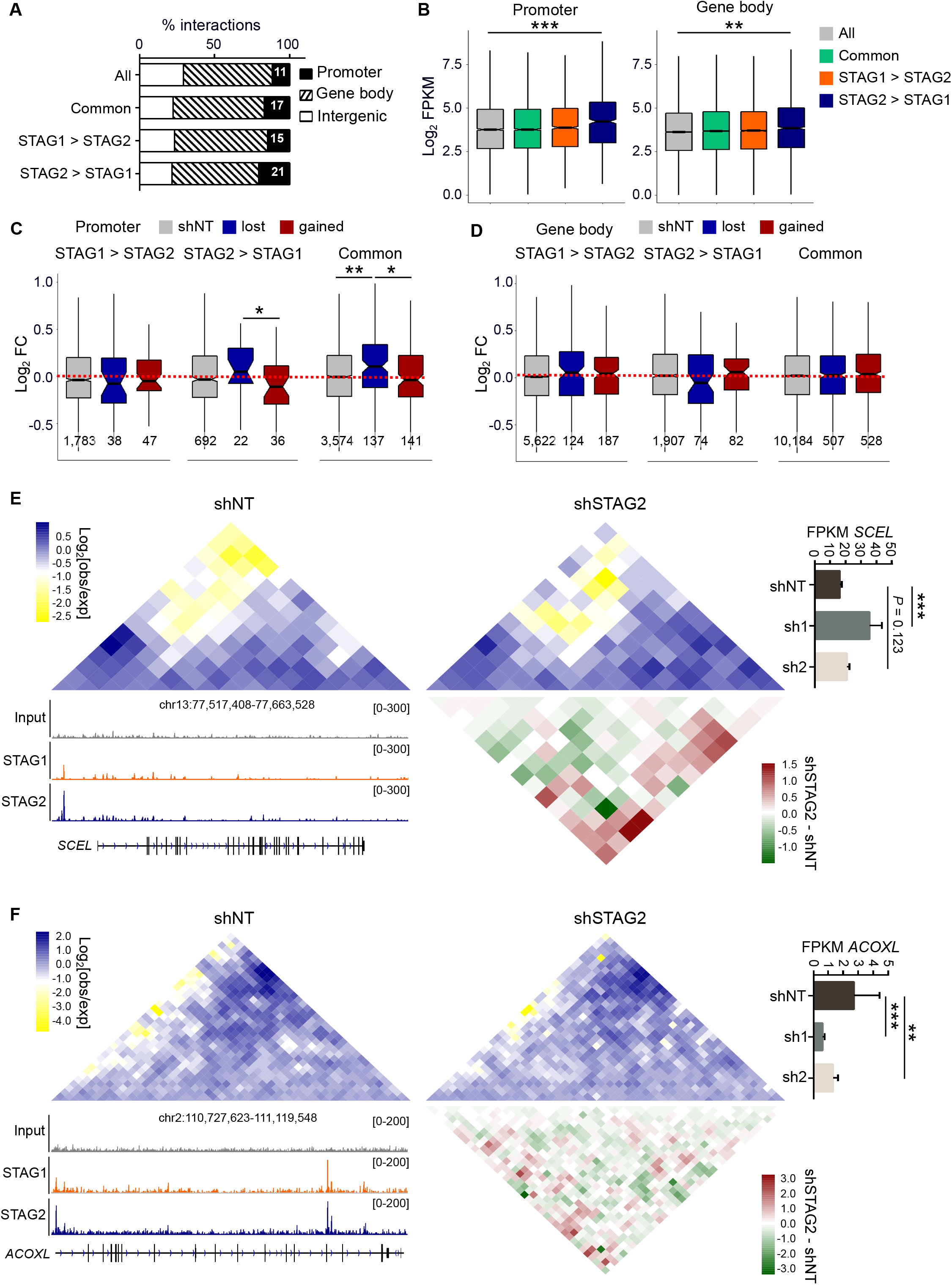
STAG2-enriched overlapping interactions engage transcriptionally active genes. (A) Distribution of all and cohesin-overlapping interactions, in control cells over genomic elements. STAG2-enriched overlapping interactions engage a higher percentage of promoters than interactions overlapping other cohesin binding sites. (B) RNA-Seq expression values (log2 FPKM) of genes engaged by all, or by cohesin-overlapping interactions, in control cells. STAG2-enriched overlapping interactions are associated with highly transcribed genes. t-test: **, *P* < 0.01; ***, *P* < 0.001. (C,D) Average fold change in gene expression values (FPKM) of genes engaged by control and differential interactions overlapping promoters (C) or gene bodies (D) and common, STAG1-enriched, or STAG2-enriched cohesin binding sites. Boxplot notches represent the confidence interval around the median. Values at the bottom of the graph refer to the number of genes engaged by every category of interactions. Mann-Whitney U test: *, *P* < 0.05; **, *P* < 0.01. (E,F) Hi-C contact matrices at the *SCEL* (E) and *ACOXL* (F) loci in control and STAG2-silenced cells. Differential contact matrices are included to emphasize the impact of STAG2 depletion on DNA contacts at these loci. Snapshots of the ChIP-Seq tracks for STAG1 and STAG2, differential contact matrices, and gene expression values (FPKM) are included. Loss of interactions overlapping the promoter of *CALD1* upon STAG2 silencing results in a consistent increase in gene expression, while gain of interactions on the promoter of *FANCE* is translated into decreased gene expression. Error bars represent mean ± SEM. t-test: **, *P* < 0.01; ***, *P* < 0.001.

### STAG2-enriched overlapping interactions engage transcriptionally active genes

To better characterize the transcriptional consequences of changes in chromatin looping, we overlapped the three subsets of interactions [control (shNT), lost, and gained] with common, STAG1-enriched, and STAG2-enriched cohesin binding sites. Cohesin-overlapping interactions showed a preference for promoters and gene bodies and, among these, those associated with STAG2-enriched cohesin target sites showed the highest enrichment in promoters (Figure 5A). Genes whose promoters/gene bodies were involved in interactions with STAG2-enriched binding sites were expressed at significantly higher levels than those engaged by interactions overlapping other cohesin binding sites (Figure 5B). Intersection of the contact and RNA-Seq data showed that loss/gain of interactions among STAG2-enriched overlapping loops had more pronounced transcriptional consequences than among STAG1-enriched or common cohesin overlapping interactions (Figure 5C,D). Intriguingly, genes associated with lost interactions were expressed at significantly higher levels than those associated to control or gained interactions (Supplementary Figure 5A-B). Loss of STAG2-enriched interactions in promoters was associated with increased gene expression (Figure 5C), whereas loss of contacts in gene bodies resulted in decreased transcription (Figure 5D). The opposite effect was observed for an increase in interaction frequency of STAG2-enriched loops. These effects were exemplified by *SCEL*, a gene associated with the basal/squamous UBC molecular subtype, and *ACOXL*, a gene that characterizes the luminal papillary UBC molecular class (Supplementary Figure 5C). Upon STAG2 silencing, *SCEL* was transcriptionally up-regulated and displayed both loss of interactions around the promoter and gain of interactions throughout the gene body (Figure 5E). In contrast, *ACOXL* was significantly down-regulated by both shRNAs and showed increased DNA contacts around the promoter and decreased interactions in the gene body (Figure 5F).

## DISCUSSION

The mechanisms whereby STAG2 acts as a tumor suppressor gene and contributes to cancer are not well established and are likely to be diverse. The lack of association between *STAG2* inactivation and aneuploidy/genomic instability in AML and UBC strongly suggests its participation through effects other than chromosome segregation (Balbás-Martínez et al. 2013; Welch et al. 2012). Recent evidence on the role of cohesin in higher-order chromatin structure and on the distinct functions of STAG1 and STAG2 in several cell types, mainly in the haematopoietic lineage, has provided support to the hypothesis that changes in gene expression may play a crucial role in the tumor suppressive role of STAG2. Yet, this notion is challenged by the fact that, in numerous cellular systems, suppression of STAG2 activity results in only modest changes at the global transcriptome level.

Importantly, these questions have not been addressed in UBC, the tumor with the highest prevalence of *STAG2* mutations. A major limitation has been the lack of adequate models. Most UBC lines are derived from muscle-invasive UBC and, therefore, few of them harbour *STAG2* mutations. Furthermore, until recently it has not been possible to permanently maintain normal urothelial cells in culture (Santos et al. 2019). Therefore, we aimed at assessing the effects of STAG2 knockdown on one of the most commonly used luminal-type UBC line: RT112. An important finding of our study is that the small fraction of the transcriptome undergoing changes upon STAG2 silencing overlaps significantly with genes that are differentially expressed in UBC with low vs. high STAG2 levels. These observations strongly support the adequacy of RT112 cells to explore the mechanisms through which STAG2 contributes to UBC.

We find that in RT112 cells - as in other cell types - a small subset of cohesin-bound sites is STAG2-enriched. These genomic sites are over-represented in promoters of active genes and their sequences are depleted of CTCF while being enriched in tissue-specific transcription factors (i.e. TFAP2A, KLF5). We also find that STAG2 depletion does not result in major changes in the genome compartmentalization and has limited effects on megabase organization, with merging of only a few adjacent TADs. However, genome interactions spanning variable distances were differently affected, with losses occurring mainly in short-range (<250 kb) contacts involving common or STAG2-enriched positions. Together with the finding that the lost interactions were statistically significantly clustered in nuclear space (as measured by their assortativity), our results indicate that loss of STAG2 impacts on genome organization at the intra-TAD level in bladder cancer cells. These structural changes appear to be functionally relevant, with genes whose promoters are associated with lost interactions being up-regulated in STAG2-depleted cells. Our findings indicate that depletion of STAG2 at gene promoters reduces repressive activity, allowing the transcriptional activation of the corresponding genes. Recently, it was reported that STAG2-cohesin promotes PRC1 recruitment and thereby contributes to Polycomb domain compaction and the formation of long-range contacts between those domains (Cuadrado et al. 2019). In our system, lost contacts after STAG2 depletion consistently span short distances and overlap with highly expressed genes, suggesting that the increase in transcriptional output does not result from de-compaction and de-repression of Polycomb targets. The discrepancies between the two cellular systems imply that STAG2-cohesin plays cell-type specific roles in the 3D organization of chromatin and may result from differential repressive effects of Polycomb on ES and differentiated cells.

Interestingly, two of the transcription factors whose binding motifs are enriched in STAG2-only positions - TFAP2A and KLF5 - show a tissue-restricted expression pattern, with high levels in squamous epithelia such as the skin and the esophagus (https://www.gtexportal.org). Activation of basal/squamous differentiation programs is a feature of a subset of UBC displaying down-regulation of luminal genes, reflecting loss-of-identity, and designated as basal/squamous-like (BASQ). There is increasing evidence that - in several epithelial tumors - the canonical vs. basal programs are regulated in a complex manner through epigenetic mechanisms and appear as a continuum rather than as dichotomous phenotypes. In bladder cancer cells, TFAP2A is repressed by PPARg, a major regulator of luminal-type tumors, and is up-regulated in BASQ-type tumors (Yamashita et al. 2019). In addition, the KLF4-driven regulon is selectively activated in BASQ tumors (Kamoun et al. 2020). Despite the association of *STAG2* mutations with papillary tumors, STAG2 knockdown resulted in the down-regulation of the luminal signature. These apparently paradoxical findings are, nevertheless, in agreement with the observation that the UBC subtype displaying the highest prevalence of *STAG2* mutations also shows a higher activation score of the basal differentiation signature than other luminal tumors (Kamoun et al. 2020). We therefore hypothesize that STAG2 plays a tumor suppressor role by establishing and/or maintaining the DNA looping required to sustain the luminal differentiation program in urothelial cells.

## METHODS

### Cell lines

RT112 bladder cancer cells used at CNIO and Institut Curie were from the same original stock; HEK293T cells (transformed human embryonic kidney) were from the ATCC. Cells were grown in Dulbecco’s Modified Eagle’s Medium (DMEM) supplemented with 10% heat-inactivated FBS (Fetal Bovine Serum) and 1 mM sodium pyruvate.

### Plasmids and lentiviral infections

Mission shRNAs (Sigma) were used for RNA interference. Two STAG2-targeting shRNAs were selected based on silencing efficacy and compared to a control non-targeting shRNA. Infectious lentiviruses were produced in HEK293T cells by FuGene-mediated transfection of the lentiviral construct together with the packaging plasmids psPAX2 and pCMV-VSVG. After transfection (48h), the medium was collected twice for an additional 48h. Viral supernatants were filtered and either frozen down in aliquots or applied on target cells in the presence of 5 mg·ml^−1^ of polybrene. Cells were harvested after 48h of puromycin selection (2 mg·ml^−1^) in serum-free medium. Gene silencing experiments were performed at high cell density and in the absence of serum to avoid cell cycle-dependent effects and to obtain homogeneous cell populations.

### Western blotting

Cell pellets were lysed in RIPA buffer supplemented with protease and phosphatase inhibitors. Following sonication, clearing by centrifugation, and quantification, proteins were subjected to SDS-PAGE. Samples were run under reducing conditions and then transferred to nitrocellulose membranes, which were blocked with TBS-Tween, 5% skim milk. Membranes were subsequently incubated with primary antibodies against STAG2 (Santa Cruz, ref. sc-81852, 1:500) or Vinculin (Sigma-Aldrich, ref. V9131-2ML, 1:2,000). After washing with TBS-Tween, membranes were incubated with horseradish peroxidase-conjugated secondary antibodies (Dako, 1:10,000) and washed. Reactions were detected using enhanced chemoluminiscence.

### ChIP sequencing for cohesin subunits and downstream analysis

ChIP-Seq was performed on RT112 in duplicates. Briefly, cells (4×10^7^) were washed with PBS, trypsinized, and resuspended in 20 mL of growing media supplemented with 1% formaldehyde for 15 min at room temperature (RT). After quenching with glycine (0.125 M final concentration), fixed cells were washed twice with PBS containing protease inhibitors, pelleted, and resuspended in lysis buffer (1% SDS, 10 mM EDTA, 50 mM Tris-HCl pH=8.1) at 2×10^7^ cells/ml. Chromatin was sonicated in a Covaris instrument for 30 min (20% duty cycle; 6% intensity; 200 cycle), yielding DNA fragments of 300-500 bp. Sonicated samples were centrifuged to pellet debris. Chromatin was quantified on a Nanodrop and a 30 μL aliquot of this material was used as input. Chromatin was diluted with buffer (1% Triton X-100, 2 mM EDTA pH=8, 150 mM NaCl, 20 mM Tris-HCl pH=8.1) supplemented with a protease inhibitor cocktail and PMSF. Samples were pre-cleared with a mix of protein A/G agarose beads (previously washed and blocked with 5% BSA) for 1 h at 4 °C. After centrifugation, supernatant was divided into aliquots of 500 *μ*g and incubated with 25 *μ*g of antibody [anti-STAG1 (source: Remeseiro et al. 2012b), anti-STAG2 (source: Remeseiro et al. 2012b), anti-SMC1 (source: Remeseiro et al. 2012b), non-related IgG]. After overnight incubation at 4°C, 100μL of pre-blocked protein A/G agarose beads were added for 2 h at 4 °C on a rotating platform to collect the immune complexes. Then, beads were sequentially washed with 1 mL of the following buffers: low-salt wash buffer (0.1% SDS, 1% Triton X-100, 2 mM EDTA, 20 mM Tris-HCl pH=8.1, 150 mM NaCl), high-salt wash buffer (0.1% SDS, 1% Triton X-100, 2 mM EDTA, 20 mM Tris-HCl pH=8.1, 500 mM NaCl), LiCl wash buffer (0.25 M LiCl, 1% NP-40, 1% deoxycholateNa, 1 mM EDTA, 10 mM Tris-HCl pH=8.1), and TE 1X (10mM Tris-HCl pH=7.5, 1mM EDTA). DNA was recovered in elution buffer (1% SDS, 0.1M NaHCO_3_) and cross-linking was reversed by overnight incubation at 65 °C. RNA and proteins were sequentially digested with 20 *μ*g of RNAse and 40 *μ*g of proteinase K. DNA was purified by phenol-chloroform extraction and resuspended in TE 0.5X.

For library preparation, 5 ng of DNA per condition were used. Samples were processed through sequential enzymatic treatments of end-repair, dA-tailing, and ligation to adapters with “NEBNext Ultra II DNA Library Prep Kit for Illumina” (New England BioLabs, ref. E7645). Adapter-ligated libraries were completed by limited-cycle PCR and extracted with a single double-sided SPRI size selection. Resulting average fragment size was 370 bp, from which 120 bp corresponded to adaptor sequences. Libraries were applied to an Illumina flow cell for cluster generation and sequenced on an Illumina NextSeq 500.

Conversion of Illumina BCL files to bam format was performed with the Illumina2bam tool (Wellcome Trust Sanger Institute - NPG). RUbioSeq (v3.8.1; Rubio-Camarillo et al. 2017) was used with default parameters to check sequencing quality, align reads to the human reference genome (hg19), normalize library sizes, and calculate ChIP-Seq peaks. Differential peaks between STAG1 and STAG2 were calculated with the DiffBind R package (Ross-Innes et al. 2012). We used the dba.count function to include peaks in the analysis that appear at least in one sample from STAG1, STAG2 or SMC1 ChIP-Seq experiments. Then, peaks were length normalized to 500bp, extending 250bp up- and down-stream of the peak summit to keep the peaks at a consistent width. Read counting in peaks was done with ChIP-Seq alignments normalized by library size. Differential enrichment peaks analyses were carried out with dba.contrast and dba.analyze functions. Three categories of cohesin-bound genomic positions were identified with dba.report function, based on statistical differences in read densities for STAG1 and STAG2: common peaks with no statistical differences between STAG1 and STAG2 read densities; STAG1-enriched (STAG1>STAG2) peaks with FDR<0.05 and higher STAG1 read density; STAG2-enriched (STAG2>STAG1) peaks with FDR<0.05 and higher STAG2 read density. Peak annotation over genomic elements was done with HOMER (v4.8.3;Heinz et al. 2010). RPKM-normalized bigwig files were generated with DeepTools (v3.0.2) bamCoverage. Heatmaps and density plots were carried out with DeepTools computeMatrix and plotHeatmap around the center of peaks. Motif enrichment was done with MEME-ChIP (v4.12.0; Machanick and Bailey 2011) using default parameters.

### ChIP-Seq for histone modifications and chromatin state assignment

ChIP-Seq for histone marks and CTCF were performed using RT112 cells. Cells were crosslinked directly in culture medium with formaldehyde (1% final concentration) for 10 min at RT. The reaction was stopped by adding Glycine (0.125 M final concentration) for 10 minutes at RT. Fixed cells were rinsed 3 times with PBS containing protease inhibitors, pelleted, and resuspended in lysis buffer (10mM EDTA, pH=8, 50mM Tris-HCl pH=8, SDS 1%). After centrifugation, ChIPs were performed using the ChIP-IT High Sensitivity kit (Active Motif, ref. 53040), following manufacturer’s instructions. Chromatin was sonicated in a Diagenode Picoruptor sonicator for 10 min (30s ON/ 30s OFF). Sheared chromatin was immunoprecipitated using the following antibodies: H3K4me1 (Abcam, ref. ab8895), H3K4me3 (Abcam, ref. ab8580), H3K27me3 (Active Motif, ref. 39155), H3K27Ac (Abcam, ref. ab4729), H3K9Ac (Millipore, ref. 07-352), H3K9me3 (Active Motif, ref. 39161), and CTCF (Millipore, ref. 07-729).

ChIP-Seq libraries were prepared using the NEXTflex ChIP-Seq Kit (Bioo Scientific, ref. 5143-02) following the manufacturer’s protocol with some modifications. Briefly, 10 ng of ChIP enriched DNA were end repaired using T4 DNA polymerase, Klenow DNA polymerase and T4 PNK, then size selected and cleaned-up using Agencourt AMPure XP beads (Beckman, ref. A63881). A single ‘A’ nucleotide was added to the 3’ ends of the blunt DNA fragments with a Klenow fragment (3’ to 5’exo minus). The ends of the DNA fragments were ligated to double stranded barcoded DNA adapters (NEXTflex ChIP-Seq Barcodes, Bioo Scientific, ref. 514120) using T4 DNA Ligase. The ligated products were enriched by PCR (2 min at 98 °C; [30 sec at 98°C, 30 sec at 65 °C, 60 sec at 72 °C] x 14 cycles; 4 min at 72 °C) and cleaned-up using Agencourt AMPure XP beads. Prior to sequencing, DNA libraries were checked for quality and quantified using a 2100 Bioanalyzer (Agilent). Libraries were loaded in the flow cell at 8pM concentration and clusters were generated using the Cbot and sequenced on the Illumina Hiseq2500 as single-end 50 base reads following Illumina’s instructions.

Sequence reads were mapped to reference genome hg19 using Bowtie 1.0.0 with the following parameters -m 1 --strata --best -y -S -l 40 -p 2. Peak detection was performed using MACS2 (model based analysis for ChIP-Seq v2.1.0.20140616) software under settings where an input sample was used as a negative control. We used a default cut-off and -B option for broad peaks.

ChromHMM was used to identify chromatin states. The genome was analysed at 200 bp intervals and the tool was used to learn models from the six histone marks, CTCF ChIP-Seq reads files and corresponding Input controls. A model of 10 states was selected and applied on all samples. 8 of the 10 states identified were then given functional annotation based on histone marks enrichment.

### RNA Sequencing and analysis

RNA-Seq of control and STAG2-silenced RT112 cells was performed in triplicates (1×10^6^ cells per sample). Total RNA was extracted using TRIzol (ThermoFisher, ref. 15596026) and purified with the RNeasy Mini Kit (Qiagen, ref. 74104), according to manufacturer’s instructions.

For library preparation, 1 μg of total RNA per condition, each containing an equal amount of ERCC ExFold RNA Spike-In Mix 2 (Ambion, ref. 4456739), was used. Average sample RNA Integrity Number was 9.4 (range 9.0 - 9.8) when assayed on an Agilent 2100 Bioanalyzer. PolyA+ fraction was purified and randomly fragmented, converted to double stranded cDNA and processed through subsequent enzymatic treatments of end-repair, dA-tailing, and ligation to adapters as in Illumina’s “TruSeq Stranded mRNA Sample Preparation Part # 15031047 Rev. D” kit (this kit incorporates dUTP during 2nd strand cDNA synthesis, which implies that only the cDNA strand generated during 1st strand synthesis is eventually sequenced). Adapter-ligated library was completed by PCR with Illumina PE primers. The resulting purified cDNA library was applied to an Illumina flow cell for cluster generation and sequenced on an Illumina HiSeq 2500.

Conversion of Illumina BCL files to bam format was performed with the Illumina2bam tool (Wellcome Trust Sanger Institute - NPG). Nextpresso 1.9 was used for downstream RNAseq analysis (Graña et al. 2017). Raw reads were aligned to the human reference genome (hg19).

### Hi-C library preparation and analysis

RT112 cells were arrested in G1 by culturing at high density and low serum (1%). Hi-C was performed as previously described (Rao et al. 2014) with some modifications. Purified DNA was fragmented to obtain fragments of an average size of 300–400 bp using a Bioruptor Pico (Diagenode; 8 cycles; 20 sec ON/60 sec OFF). 3 μg of DNA per condition were used for library preparation. Biotinylated DNA was pulled down with Dynabeads MyOne T1 streptavidin beads. End-repair, A-tailing and the Illumina adaptors ligation were performed on beads. Libraries were amplified by 10 cycles of PCR and purified using AMPure XP beads. The concentration and size distribution of the Hi-C library after PCR amplification were determined using a Qbit fluorometer and visual exploration in an agarose gel. Hi-C libraries were then paired-end sequenced on an Illumina NextSeq500 (200M reads per library).

Data were processed for read quality control, mapping, interaction detection, filtering, and matrix normalization using TADbit (Serra et al. 2017) (Supplementary Figure 6A). First, reads were quality-controlled using the TADbit implementation of FastQC for Hi-C datasets. Average PHRED scores were >25 throughout paired-end reads, indicative of good quality (Supplementary Figure 6B). Then, we used a fragment-based strategy for mapping the remaining reads to the reference human genome (GRCh38). Non-informative contacts including self-circles, dangling-ends, errors, random breaks or duplicates were filtered out, resulting in 158-197M valid interactions per condition (Supplementary Table 1). These were then used to generate genome-wide interaction maps at 100kb and 20kb resolution to segment the genome into A/B compartments, demarcate TADs, and identify changes in chromatin looping. A/B compartments were identified with HOMER (Heinz et al. 2010) by calculating the first two eigenvectors of vanilla-normalized 100 kb contact matrices for every chromosome. Chromosome bins with positive PC1 values and high gene density were considered to be part of compartment A, while bins with negative values and low gene density were assigned to compartment B. For chromosome 4, there was no clear separation between the first and second eigenvector profiles, so the values of the second eigenvector were taken into account for further analysis. The Y chromosome was excluded from the analysis of genomic bins switching compartments.

TADs were identified with the TAD detection algorithm implemented in TADbit using vanilla-normalized 100kb contact matrices. TAD border localization and strength were calculated to evaluate their conservation upon depletion of STAG2. Finally, 20kb matrices were used to identify significant interactions in control (shNT) and STAG2 depleted cells with HOMER analyzeHiC (FDR < 0.1) (Heinz et al. 2010). HOMER analyzeHiC was also employed to detect differential interactions between shNT and STAG2-silenced cells. Only high scoring differential interactions (*P* < 0.05) in both *STAG2*-targeting shRNAs were retained for downstream analyses.

### Assortativity of regions altered upon STAG2 silencing

To assess whether the chromatin contact regions lost or gained upon STAG2 silencing, or the regions coding for genes deregulated upon STAG2 silencing, are located in proximity in 3D, chromatin assortativity (ChAs) analysis was used (Pancaldi et al. 2016). We represent the genome as a network of nodes (chromatin fragments, here corresponding to Hi-C bins) which are connected if a Hi-C contact between them is observed. Networks were displayed using Cytoscape organic layout.

To select significant Hi-C contacts we identified significant Hi-C interactions at specific binning resolution (20kb, 100kb, 1mb) by comparing each dataset to the background of the same datasets. Each sample is considered as independent. We can thus define a control network and compare it to the STAG2 silenced network and identify connections that are lost and gained from one condition to the next. Briefly, ChAs is a measure of correlation of feature values across all edge pairs in a network and allows us to see whether nodes with a specific property tend to interact more with each other than expected at random. We asked whether assortativity of chromatin regions affected by STAG2 removal is particularly strong for contacts spanning specific ranges of genomic distance by filtering the contacts by distance spanned, eliminating contacts spanning progressively longer distances, thus generating networks in which the minimum distance spanned by any contact is 50kb, 100kb, 250kb, 500kb, 750kb, 1Mb. Finally we mapped network nodes to genes by finding which Hi-C bins were overlapping promoters and thus filtered ‘in-silico’ promoter-promoter networks from our Hi-C networks at the different resolutions. We then assigned to all the nodes in the P-P network the value of Log2FC of the gene between STAG2 KO and WT and ChAs of fold change. We repeated the calculation of assortativity in networks on which we permuted the expression values on the network nodes to generate a null distribution of ChAs values and show the distribution of these random ChAs values in the plots.

## DATA ACCESS

The following secure token has been created to allow review of record GSE111913 encompassing the ChIP-Seq data on cohesin subunits while it remains in private status: gdstsouqtdendir. ChIP-Seq data on histone modifications and CTCF for RT112 are deposited in the GEO database under accession number GSE104804. RNA-Seq data for RT112 cells next-generation sequencing data are available in NCBI GEO, under accession number GSE154878. Hi-C data are available in NCBI GEO, under accession number GSE155380. The authors declare that all other data supporting the findings of this study are within the manuscript and its supplementary files.

## ACKNOWLEDGMENTS

We thank all the staff of the GenomEast high throughput sequencing facility of the IGBMC in Strasbourg (CNRS UMR7104, Inserm U1258, Université de Strasbourg) for the sequencing of ChIPs of histone marks and CTCF; the UROMOL investigators and Lars Dyrskjot for providing RNA-Seq data; Ana Losada for providing reagents; and Mark Kalisz, Ana Losada, and Ana Cuadrado for critical review of the manuscript.

## AUTHOR CONTRIBUTIONS

LR, EL, MAM-R, and FXR initiated and designed the studies. LR and YC performed the Hi-C experiment and LR analysed and integrated it with the RNA-Seq and ChIP-Seq data. EL and MC performed RNA-Seq and cohesin ChIP-Seq experiments and ECP, EL, and LR analysed the data. VP, MMM and JAR performed the assortativity analyses and helped with the comparison of the RNA-Seq data on RT112 cells and the human tumor data. HNK and FR conducted the ChIP-Seq experiments for histone modifications and CTCF and the analysis of chromatin states in RT112 cells. FLD supervised and provided guidance on the preparation of the Hi-C libraries. FS supervised and provided guidance on the analysis of the Hi-C data. LR, EL, VP, MAM-R, and FXR interpreted the results and wrote the manuscript. MAM-R and FXR supervised the whole research and provided guidance. All authors had access to the final manuscript and approved the submission of the final version of the paper.

## FIGURE LEGENDS

**Supplementary Figure 1.**
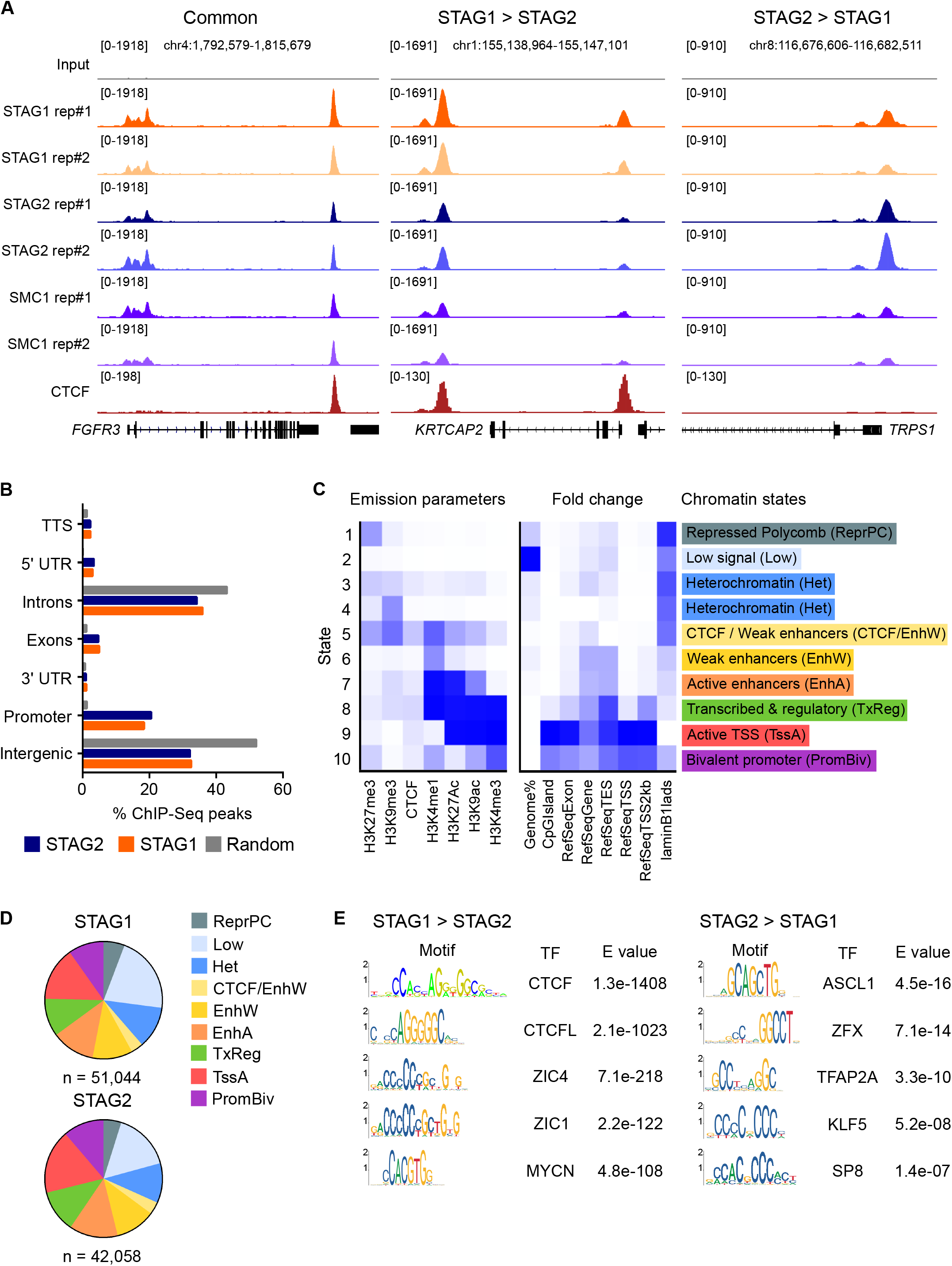
STAG1- and STAG2-enriched cohesin sites show differential overlap with the insulator CTCF. (A) Snapshots of STAG1, STAG2, and CTCF ChIP-Seq tracks at loci representative of the three categories of cohesin-bound genomic sites. (B) Bar-plot diagram showing distribution of STAG1 and STAG2 binding sites over genomic elements. (C) Chromatin state assignments for RT112 cells made by ChromHMM. The left panel displays a heatmap of the emission parameters in which each row corresponds to a different state and each column corresponds to a different histone modification/CTCF. The darker blue reflects a greater probability of observing the mark in the state. The heatmap to the right displays the fold enrichment for external genomic annotations at each chromatin state. (D) Distribution of STAG1 and STAG2 target sites over chromatin states defined in (C). (E) Motif enrichment analysis of STAG1- and STAG2-enriched positions. E values of the top 5 enriched transcription factor motifs are shown.

**Supplementary Figure 2.**
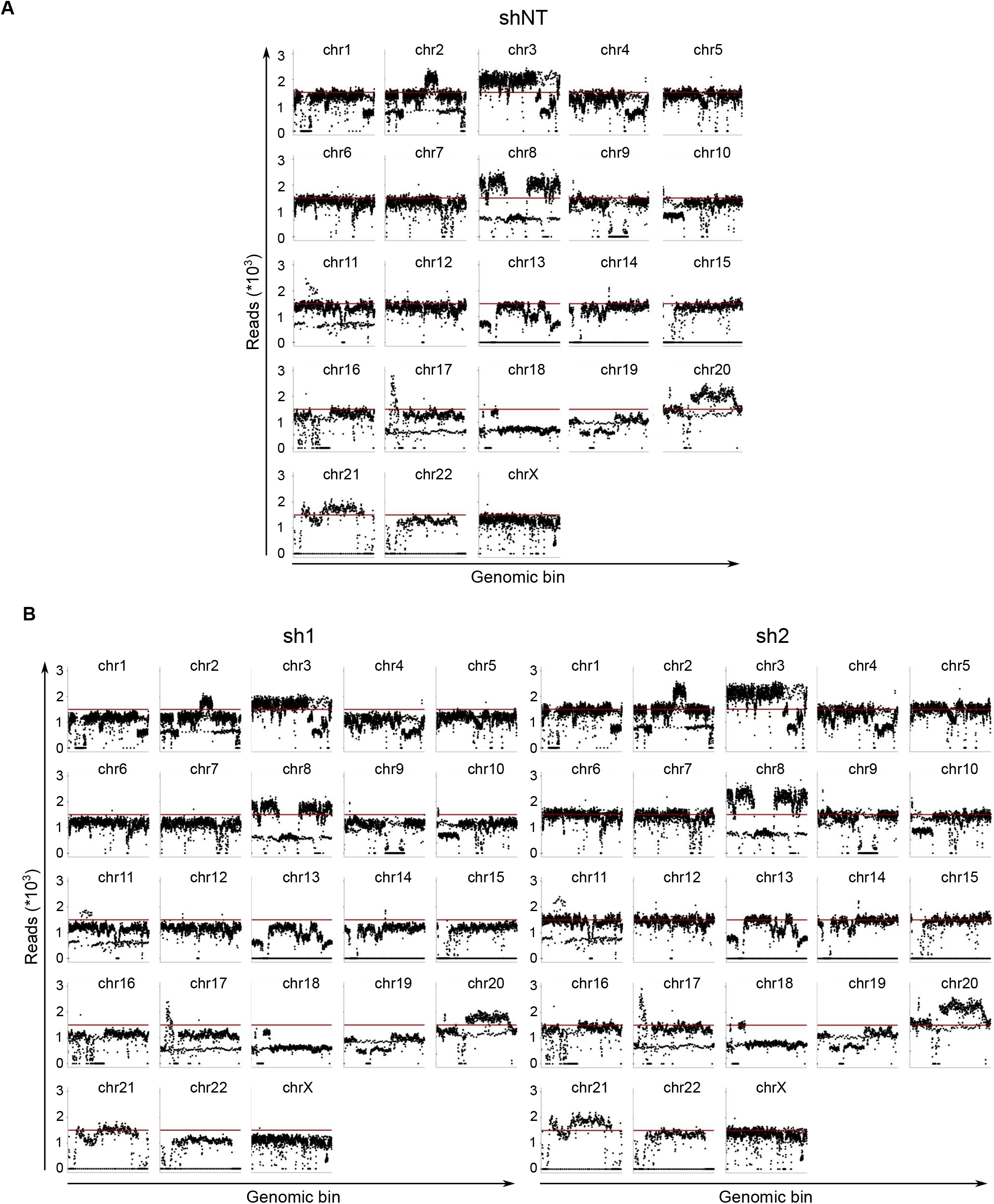
STAG2 silencing in RT112 cells does not result in major changes in gene copy number. (A,B) Read counts per 100 kb-sized genomic bin along each chromosome in control (A) and STAG2-silenced RT112 cells (B).

**Supplementary Figure 3.**
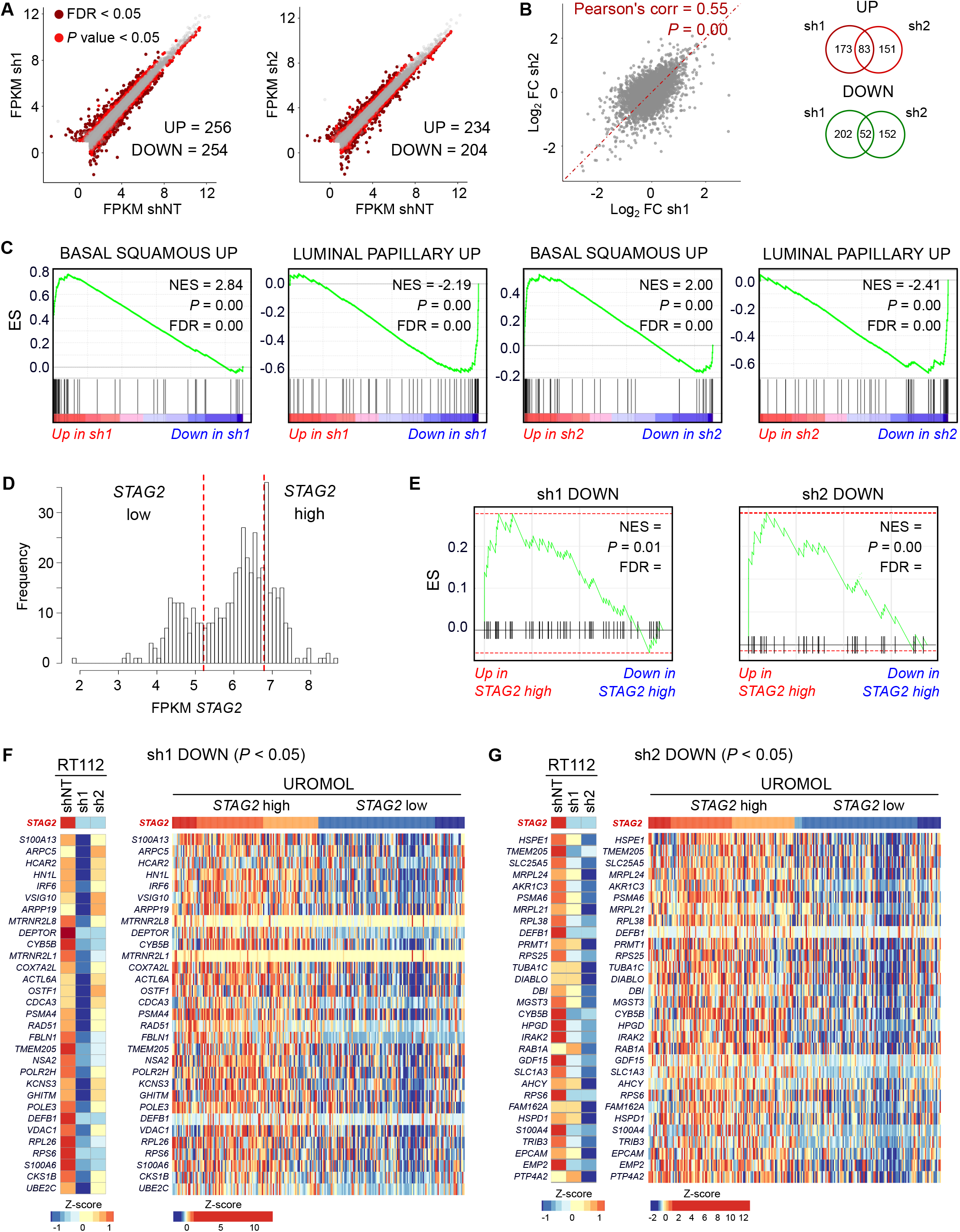
Gene expression changes upon STAG2 silencing in RT112 cells. (A) Scatter plots of expression values (FPKM) of genes in control versus STAG2-silenced cells. Statistically significant differentially expressed genes are highlighted in dark (FDR < 0.05) or light red (*P* < 0.05). (B) Scatter plot showing a positive and significant correlation between gene expression changes in sh1 and sh2 (left). Venn diagrams displaying the overlap between sh1 and sh2 in terms of significant up- and down-regulated genes. (C) GSEA enrichment plots of gene sets associated with the luminal and basal subtypes of muscle-invasive UBC showing significant deregulation in STAG2-silenced RT112 cells. (D) Distribution of *STAG2* expression (FPKM) in the UROMOL cohort of 476 UBC samples (Hedegaard et al. 2016), highlighting the thresholds of the first and fourth quartiles (119 samples per group). We defined “*STAG2* high” cases as those with expression values in the fourth quartile, and “*STAG2* low” cases as those with *STAG2* levels in the first quartile. (E) GSEA enrichment plots for genes down-regulated in STAG2-silenced cells in “*STAG2* high” versus “*STAG2* low” tumor samples. (F,G) Heatmaps displaying relative expression values (Z-score of FPKM) of genes significantly down-regulated in RT112 cells with sh1 (F) or sh2 (G) and in “*STAG2* low” versus “*STAG2* high” tumor samples.

**Supplementary Figure 4.**
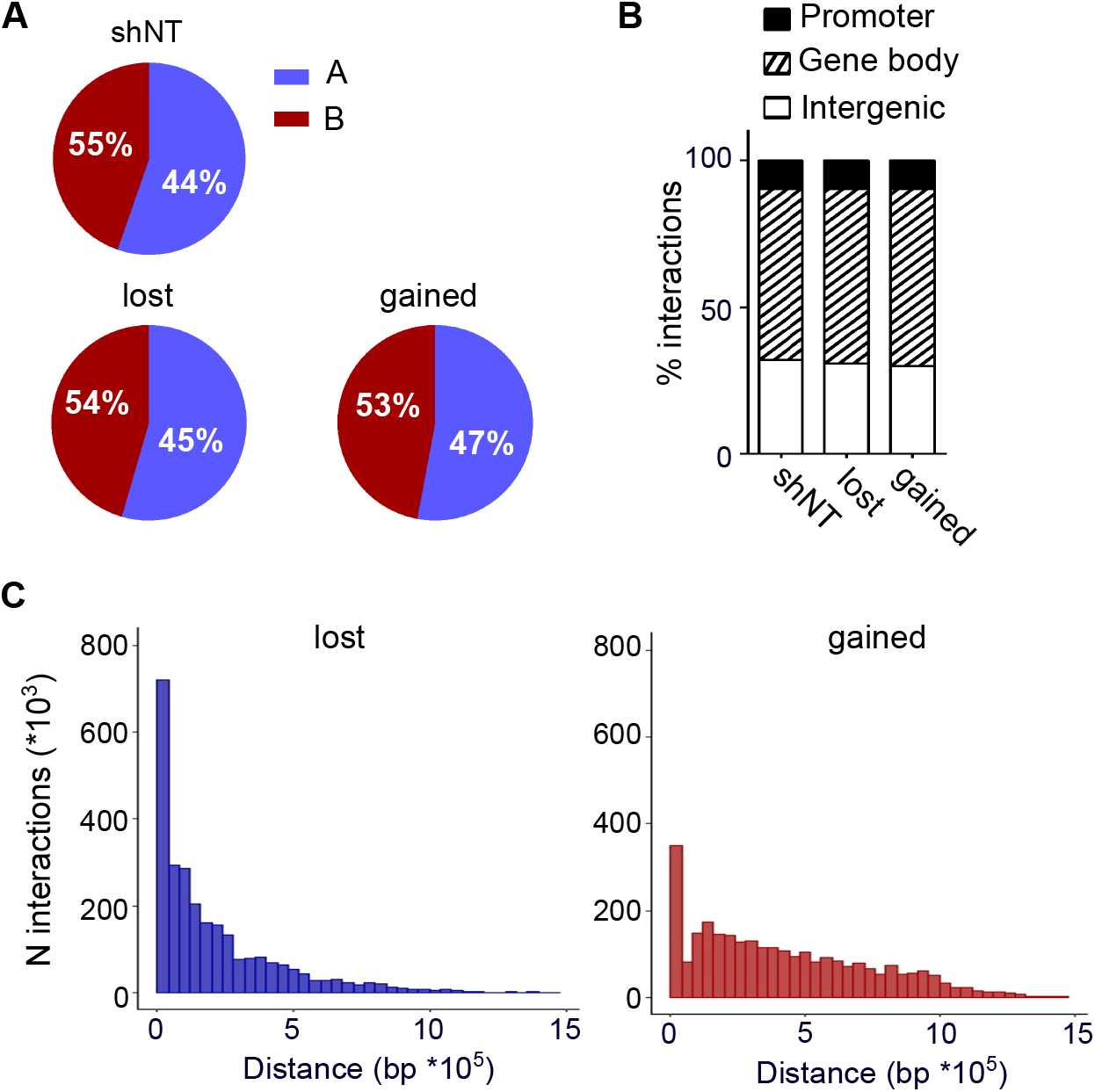
Genomic characterization of lost and gained interactions in RT112 cells upon STAG2 knockdown. (A,B) Overlap of control (shNT), lost, and gained interactions with A/B compartments (A) and genomic elements (B). (C) Distribution of distances spanned by lost and gained interactions.

**Supplementary Figure 5.**
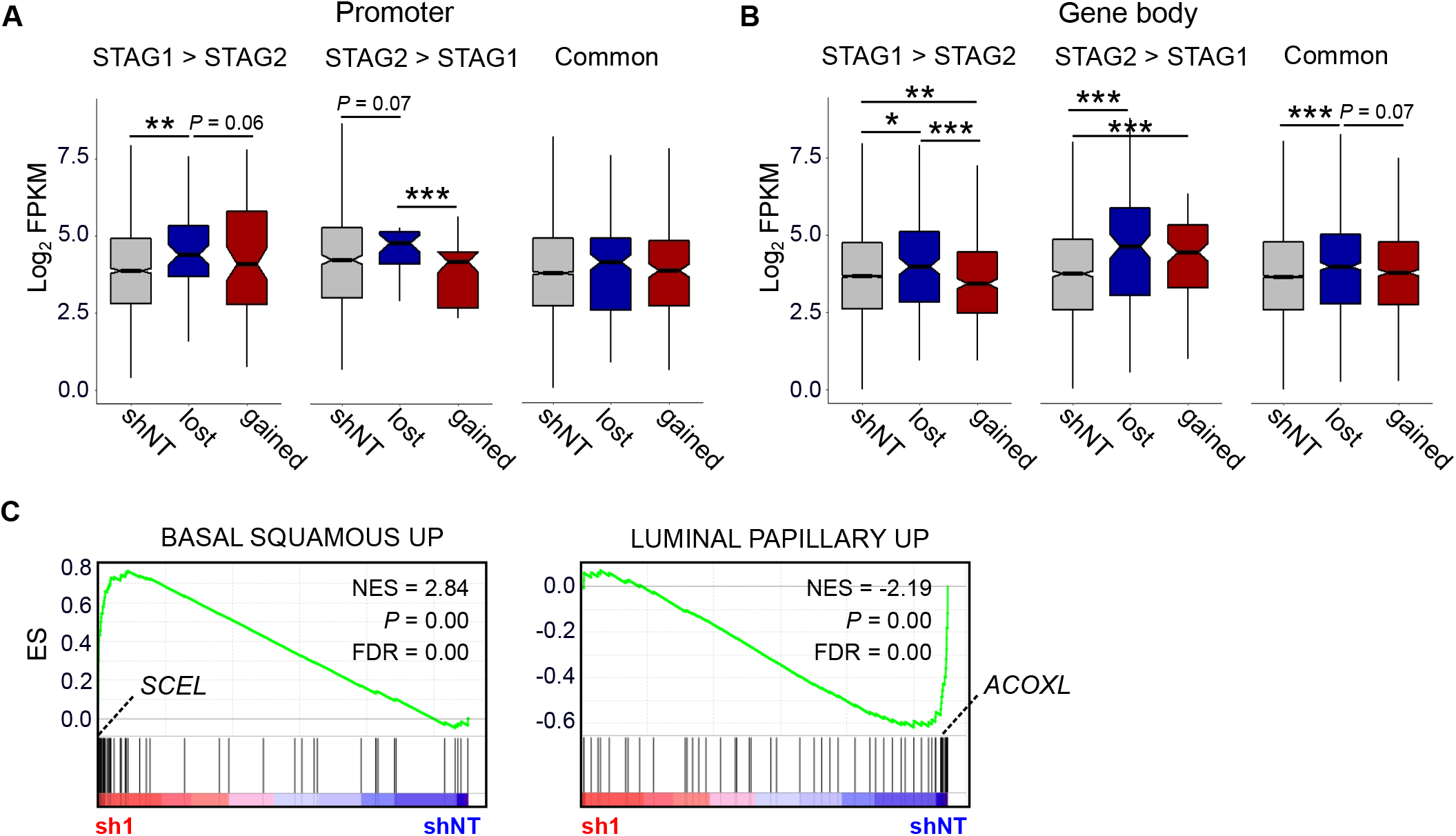
Analysis of the consequences of changes in DNA looping on gene expression. (A,B) Expression values of genes engaged by control and differential interactions overlapping promoters (A) or gene bodies (B) and common, STAG1-enriched, or STAG2-enriched cohesin binding sites. Boxplot notches represent the confidence interval around the median. (C) Gene expression values (FPKM) of *SCEL* and *ACOXL* in control and STAG2-silenced cells.

**Supplementary Figure 6.**
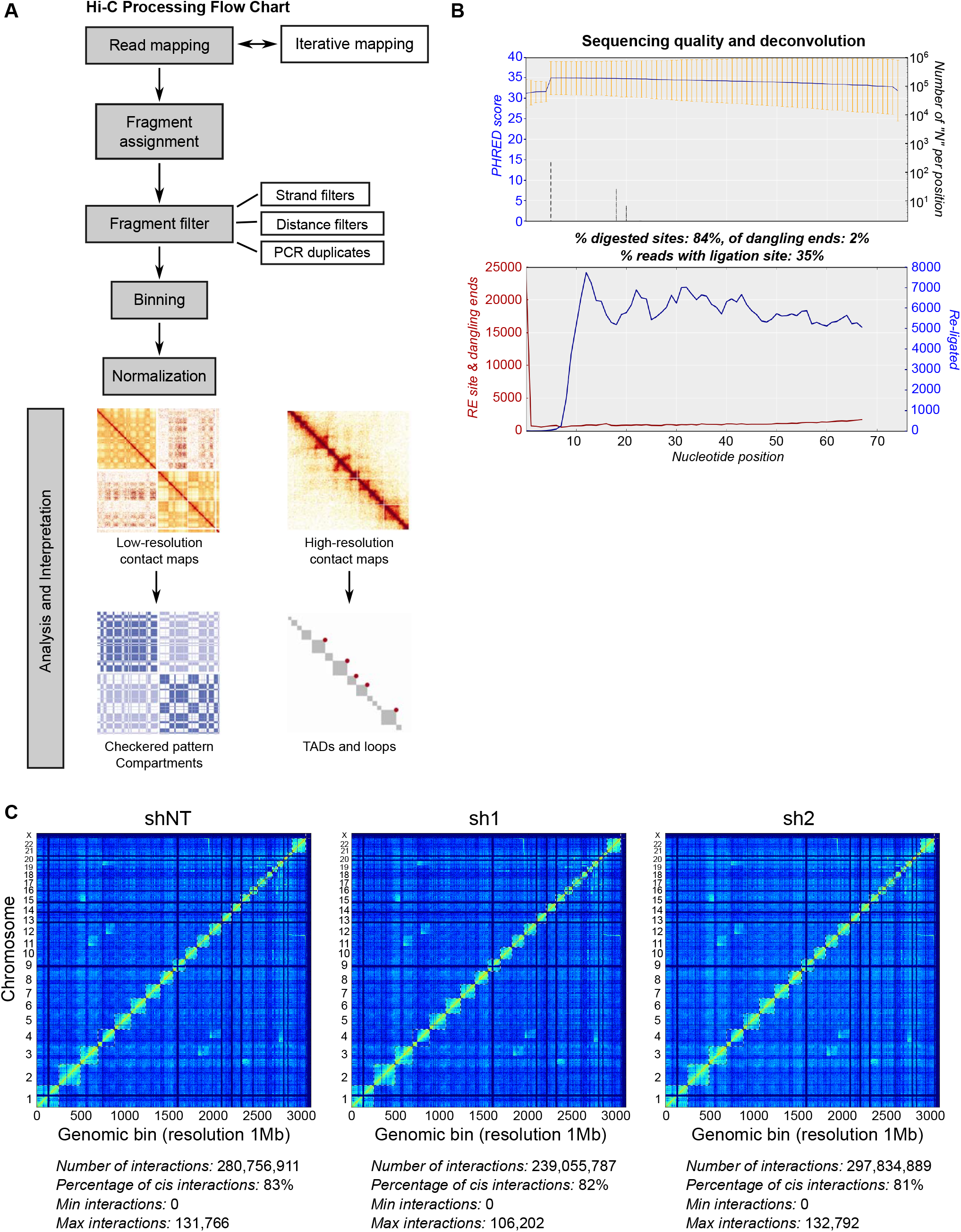
Quality check Hi-C datasets. (A) Pipeline for the analysis of Hi-C data. (B) Example of the quality report on the shNT read 1 FASTQ file created by the TADbit pipeline. The report includes information on the efficiency of both digestion and ligation. (C) Filtered genome-wide Hi-C interaction maps at 1Mb resolution for each sample type. Pairs of loci that reside on the same chromosome show higher interaction frequencies than loci that reside on different chromosomes.

**Supplementary Table 1.** Number of valid read pairs per sample after filtering.

| **Name** | **Counts** |  |  |
| --- | --- | --- | --- |
|  | *shNT* | *sh1* | *sh2* |
| self-circle | 406,110 | 261,567 | 851,346 |
| dangling-end | 16,128,197 | 14,301,386 | 31,266,244 |
| error | 1,531,788 | 1,037,286 | 2,219,148 |
| extra dangling-end | 50,214,145 | 43,467,655 | 101,931,934 |
| too close from RES | 71,519,086 | 61,189,441 | 146,539,702 |
| too short | 12,682,490 | 10,843,032 | 25,512,686 |
| too large | 5,347 | 3,638 | 10,436 |
| over-represented | 3,094,311 | 2,514,162 | 5,977,962 |
| duplicated | 47,447,543 | 46,562,136 | 323,326,328 |
| random breaks | 340,393 | 244,788 | 696,764 |
| valid-pairs | 183,956,218 | 158,263,139 | 197,459,063 |

